# Signatures of long-term balancing selection in human genomes

**DOI:** 10.1101/119529

**Authors:** Bárbara Domingues Bitarello, Cesare de Filippo, João Carlos Teixeira, Joshua M. Schmidt, Philip Kleinert, Diogo Meyer, Aida M. Andrés

**Affiliations:** Department of Genetics and Evolutionary Biology, University of São Paulo, São Paulo; Evolutionary Genetics Department, Max Planck Institute for Evolutionary Anthropology, Leipzig; Unit of Human Evolutionary Genetics, Institut Pasteur, Paris; Current address: Computational Molecular Biology Department, Max Planck Institute for Molecular Genetics, Berlin; UCL Genetics Institute, Department of Genetics, Evolution and Environment, University College London, London

**Keywords:** natural selection, overdominance, site frequency spectrum, neutrality test, summary statistic, genome-wide scan

## Abstract

Balancing selection maintains advantageous diversity in populations through various mechanisms. While extensively explored from a theoretical perspective, an empirical understanding of its prevalence and targets lags behind our knowledge of positive selection. Here we describe the *Non-Central Deviation* (*NCD*), a simple yet powerful statistic to detect long-term balancing selection (LTBS) that quantifies how close frequencies are to expectations under LTBS, and provides the basis for a neutrality test. *NCD* can be applied to a single locus or genomic data, and can be implemented considering only polymorphisms (*NCD1*) or also considering fixed differences with respect to an outgroup (*NCD2*) species. Incorporating fixed differences improves power, and *NCD2* has higher power to detect LTBS in humans under different frequencies of the balanced allele(s) than other available methods. Applied to genome-wide data from African and European human populations, in both cases using chimpanzee as an outgroup, *NCD2* shows that, albeit not prevalent, LTBS affects a sizable portion of the genome: about 0.6% of analyzed genomic windows and 0.8% of analyzed positions. Significant windows (*p* < 0.0001) contain 1.6% of SNPs in the genome, which disproportionally fall within exons and change protein sequence, but are not enriched in putatively regulatory sites. These windows overlap about 8% of the protein-coding genes, and these have larger number of transcripts than expected by chance even after controlling for gene length. Our catalog includes known targets of LTBS but a majority of them (90%) are novel. As expected, immune-related genes are among those with the strongest signatures, although most candidates are involved in other biological functions, suggesting that LTBS potentially influences diverse human phenotypes.

## Introduction

Balancing selection refers to a class of selective mechanisms that maintains advantageous genetic diversity in populations. Decades of research have established HLA genes as a prime example of balancing selection (Meyer & Thomson 2001; Spurgin & Richardson 2010), with thousands of alleles segregating in humans, extensive support for functional effects of these polymorphisms (e.g. Prugnolle et al. 2005), and various well-documented cases of association between selected alleles and disease susceptibility ( e.g. Raychaudhuri et al. 2012; Howell 2014). The catalog of well-understood non-HLA targets of balancing selection in humans remains small, but includes genes associated to phenotypes such as auto-immune diseases (Ferrer-Admetlla et al. 2008; Sironi & Clerici 2010), resistance to malaria (Malaria Genomic Epidemiology Network 2015), HIV infection (Biasin et al. 2007) or susceptibility to polycystic ovary syndrome (Day et al. 2015). Thus, besides historically influencing individual fitness, balanced polymorphisms shape current phenotypic diversity and susceptibility to disease.

Balancing selection encompasses several mechanisms (reviewed in Andrés 2011; Key, Teixeira, et al. 2014; Fijarczyk and Babik 2015). These include heterozygote advantage or overdominance, some cases of frequency-dependent selection (e.g. rare allele advantage), selective pressures that fluctuate in time (e.g. Bergland et al. 2014) or in space in panmitic populations (Charlesworth et al. 1997), and some cases of pleiotropy (Johnston et al. 2013). For overdominance, pleiotropy, and some instances of spatially variable selection, a stable equilibrium - at the frequency equilibrium, which maximizes fitness in the population - can be reached (Charlesworth & Charlesworth 2010). For other mechanisms, the frequency of the selected allele can change in time without reaching a stable equilibrium. Regardless of the mechanism, long-term balancing selection (LTBS) has the potential to leave identifiable signatures in genomic data. The first is an increase in the ratio of polymorphic to divergent sites: by reducing the probability of fixation of a variant, balancing selection increases the local time to the most recent common ancestor (Hudson & Kaplan 1988) and the density of polymorphisms. The HKA test is commonly used to detect this signature (Hudson et al. 1987). The second signature is an excess of alleles segregating at intermediate frequencies. In humans, the folded site frequency spectrum (SFS) – the frequency distribution of minor allele frequencies (MAF) — is typically L-shaped, showing an excess of low-frequency alleles when compared to expectations under neutrality and demographic equilibrium. Regions under LTBS, on the other hand, can show a markedly different SFS, with proportionally more alleles at intermediate frequency. Such a deviation in the SFS is identified by classical neutrality tests such as Tajima’s *D* (Taj*D*) and newer statistics such as MWU-high (Nielsen et al. 2009). In some cases, very ancient balancing selection can maintain trans-species polymorphisms in sister species (Leffler et al. 2013; Teixeira et al. 2015). On the other hand, when balancing selection is very recent or transient (Sellis et al. 2011), signatures are difficult to distinguish from incomplete, recent selective sweeps (Key, Teixeira, et al. 2014).

While balancing selection has been extensively explored from a theoretical perspective, an empirical understanding of its prevalence lags behind our knowledge of positive selection. This stems from technical difficulties in detecting balancing selection, as well as the perception that it may be a rare selective process. In fact, few methods have been developed to identify its targets, and only a handful of studies have sought to uncover them genome-wide in humans (Asthana et al. 2005; Bubb et al. 2006; Alonso et al. 2008; Andrés et al. 2009; Leffler et al. 2013; DeGiorgio et al. 2014; Rasmussen et al. 2014; Siewert & Voight 2017). Some studies have identified genes (Andrés et al. 2009) or genomic regions (DeGiorgio et al. 2014) with an excess of polymorphisms and intermediate frequency alleles, while others have identified trans-species polymorphisms between humans and their closest living relatives (chimpanzees and bonobos) (Leffler et al. 2013; Teixeira et al. 2015). Overall, these studies suggested that balancing selection may act on a small portion of the genome, although the limited extent of data used (e.g. exome data (Andrés et al. 2009), small sample size (DeGiorgio et al. 2014)), and stringency of the criteria (e.g. trans-species polymorphisms predating human-chimpanzee divergence (Leffler et al. 2013; Teixeira et al. 2015)) may underlie the paucity of detected regions.

In the tradition of neutrality tests analyzing the SFS directly (e.g. Nielsen et al. 2005; Williamson et al. 2007; Nielsen et al. 2009), we propose the “Non-Central Deviation” (*NCD*) statistic, which measures the degree to which the local SFS deviates from expectations under balancing selection using only polymorphisms (*NCD1*) or also fixed differences to an outgroup (*NCD2*). We show, through simulations, that *NCD2* outperforms existing methods under a realistic demographic scenario for human populations. We apply *NCD2* to genome-wide data from four human populations to test the null hypothesis of neutral evolution, and use both outlier and simulation-based approaches to identify genomic regions bearing signatures of LTBS.

## Methods

### The Non-Central Deviation (NCD) statistic

#### Background

Owing to linkage, the signature of long-term balancing selection extends to the genetic neighborhood of the selected variant(s); therefore, patterns of polymorphism and divergence in a genomic region can be used to infer whether it evolved under LTBS (Charlesworth 2006; Andrés 2011). As mentioned, LTBS leaves two distinctive signatures in linked variation, when compared with neutral expectations: an increase in the ratio of polymorphic to divergent sites and an excess of alleles segregating at intermediate frequencies when compared to neutral expectations (Figure 1A-B).With heterozygote advantage, the frequency equilibrium (*f*_eq_) depends on the relative fitness of each genotype (Charlesworth & Charlesworth 2010): when the two types of homozygotes have the same fitness (symmetric overdominance), *f*_eq_ = 0.5; when the fitness of the two homozygotes is different (asymmetric overdominance), *f*_eq_ ≠ 0.5. Under negative frequency-dependent selection and fluctuating selection, while an equilibrium may not be reached (suppl. information S1), *f*_eq_ can be thought of as the frequency of the balanced polymorphism at the time of sampling. We focus on overdominance, as comparing different mechanisms of balancing selection falls outside of the scope of the paper.

**Fig. 1.**
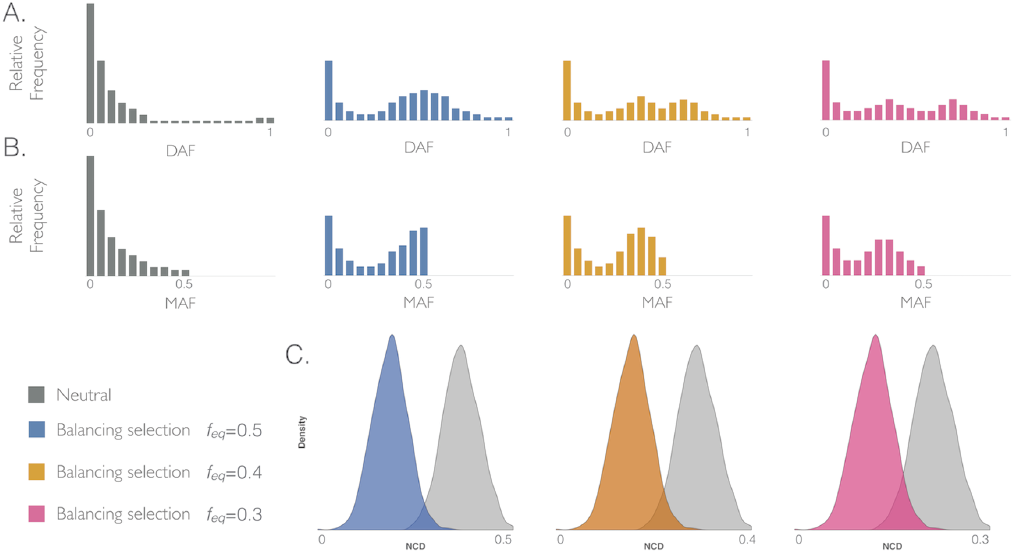
A schematic representation of site frequency spectra (SFS) under neutrality and selection, which motivates the NCD statistic. (**A**) Unfolded SFS (ranging from 0 to 1) of derived allele frequencies (DAF) for loci under neutrality (grey) or containing one site under balancing selection with frequency equilibrium (*f*_eq_) of 0.5 (blue), 0.4 (orange) and 0.3 (pink). (**B**) Folded SFS (ranging from 0 to 0.5) for minor allele frequencies (MAF). Colors as in **A**. (**C**) Distribution of *NCD* (Non-Central Deviation) expected under neutrality (grey) and under selection assuming *tf* = *f*_eq_. Colors as in **A.** *x*-axis shows minimum and maximum values that *NCD* can have for a given *tf*.

Under a model of balancing selection, we define *tf* as the expected frequency of a balanced allele, with the *NCD* statistic quantifying how far the sampled SNP frequencies are from it. Because bi-allelic loci have complementary allele frequencies, and there is no prior expectation regarding whether ancestral or derived alleles should be maintained at higher frequency, we use the folded SFS (Figure 1B). *NCD* is defined as:

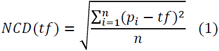

where *i* is the *i-*th informative site in a locus*, pi* is the MAF for the *i-th* informative site, *n* is the number of informative sites, and *tf* is the target frequency with respect to which the deviations of the observed alleles frequencies are computed. Thus, *NCD* is a type of standard deviation that quantifies the dispersion of allelic frequencies from *tf*, rather than from the mean of the distribution. We note that *tf* is not an estimated quantity, but rather a hypothetical frequency expected under balancing selection, with respect to which we quantify deviations in genomic data. We consider the same *tf* for all SNPs in a given locus/window because we expect most small regions to have only one balanced allele, with the rest being shifted close to the *tf* (though often not reaching it) by linkage. Low *NCD* values reflect a low deviation of the SFS from a pre-defined *tf* (Figure 1C), as expected under LTBS (suppl. information S1 and fig. S1). Of course, a priori *f*_eq_ is unknown, and thus the choice of a *tf* value may not be straightforward. We show that *NCD* values correlate strongly across *tf* values (see below), so while this is a key value in the statistic, it has limited influence in its applications. In any case, we propose below a practical approach to deal with the inherent *f*_eq_ uncertainty.

We propose two *NCD* implementations: *NCD1* uses only polymorphic sites, and *NCD2* also includes the number of fixed differences (FDs) relative to an outgroup species (i.e, all informative sites, ISs = SNPs + FDs, are used to compute the statistic). In *NCD2*, FDs are considered informative sites (ISs) with MAF = 0; thus, the greater the number of FDs, the larger the *NCD2* and the weaker the support for LTBS (suppl. fig. S2). Because *NCD2* includes also fixed differences, it uses more information and we focus on it. Nevertheless, *NCD1* can be used in the absence of an outgroup, and we show its power statistics in detail in the supplements (suppl. information S1 and suppl. table S1).

### Simulations and power analyses

*NCD*’s power was evaluated by simulations (under neutrality and with selection) with MSMS (Ewing & Hermisson 2010) following the demographic model described in Gravel et al. (2011) for African, European, and East Asian human populations, and considering a generation time of 25 years, mutation rate of 2.5 × 10^−8^ per site, recombination rate of 1 × 10^−8^, and the human-chimpanzee split at 6.5 mya. For simulations with selection, a balanced polymorphism was added to the center of the simulated sequence and modeled to achieve a pre-specified frequency equilibrium (*f*_eq_ = 0.3, 0.4, 0.5) following an overdominant model (suppl. information S1). Simulations with and without selection were run for different sequence lengths (*L* = 3, 6, 12 kb) and times of onset of balancing selection (1, 3, 5 mya). For each combination of parameter, 1,000 simulations were used to compare the relationship between true (TPR, the power of the statistic) and false (FPR) positive rates for the NCD statistics, with a FPR set to 0.05. When comparing performance under a given condition, power was averaged across demographic scenarios, *L*, and *Tbs*.

When comparing *NCD* performance to classical methods (Taj*D,* HKA), we used *NCD* optimal conditions: 3 kb, *Tbs* = 5 mya (see Results, Figure 2), which is also optimal for these other tests (suppl. table S1). For the ß statistic (Siewert & Voight 2017) we used the folded version of the test and, for consistency, also of 3 kb windows (but we note that results do not change with 1 kb windows; suppl. table S1 Table and information S1). For the *T*_1_ and *T*_2_ statistics, which require longer genomic regions to identify the signature of balancing selection, power was reported based on windows of 100 informative sites up and downstream of the target site, following BALLET’s original publication (DeGiorgio et al. 2014). Power values are for a sample size of 50 diploid individuals, but sample sizes of 30 and 10 individuals were also explored (suppl. information S1).

**Fig. 2.**
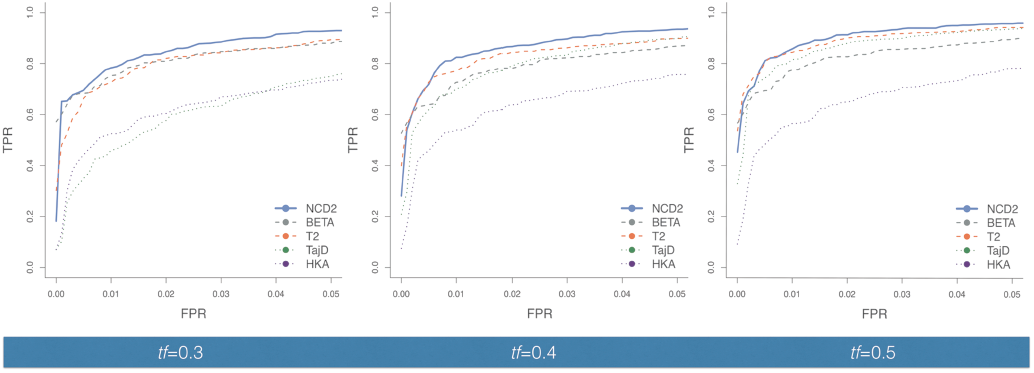
Power to detect balancing selection for *NCD2 (0. 5)* and other tests. The ROC curves summarize the true positive rate (TPR) as a function of the false positive rate (FPR) to detect LTBS for simulations where the balanced polymorphism was modeled to achieve *f*_eq_ of **(A)** 0.3, **(B)** 0.4, and **(C)** 0.5. Plotted values are for the African demography, *Tbs* = 5 mya. *L* = 3 kb, except for *T*_1_ and *T*_2_ where *L* = 100 ISs (see Methods). BETA refers to the ß statistic (Siewert & Voight 2017). For *NCD2* calculations, *tf* = *f*_eq_. European demography yields similar results (suppl. fig. S10). Power for *NCD1, NCD1+HKA* and *T1* is provided in S1 Table.

### Identifying signatures of LTBS in human genomes

#### Statistic

The implementation of the genome-wide analysis was informed by the power analyses (see Results). As our power results show, *NCD2* has very high power. While the *f*_eq_ of a putatively balanced allele -- i.e, the actual frequency of the balanced polymorphism at the time of sampling – is unknown, the simplicity of the *NCD* statistics makes it trivial to run for several *tf* values, allowing detection of balancing selection for a range of equilibrium frequencies. For each window in each population we thus calculated *NCD2* for three *tf* values (0.3, 0,4, 0.5) although in neutral simulations these results are highly correlated (suppl. fig. S9).

#### Data

We analyzed genome-wide data from two African (YRI: Yoruba in Ibadan, Nigeria; LWK: Luhya in Webuye, Kenya) and two European populations (GBR: British, England and Scotland; TSI: Toscani, Italy) from the 1000 Genomes (1000G) Project phase I (Abecasis et al. 2012), excluding SNPs only detected in the high coverage exome sequencing to avoid SNP density differences between coding and non-coding regions. We randomly sampled 50 unrelated individuals from each population (as in Key, Peter, et al. 2014) and did not consider Asian populations due to lower *NCD* performance for these populations according to our simulations (suppl. table S1, suppl. figs. S7-8). We only included in our analyses positions that passed mappability (50mer CRG, 2 mismatches (Derrien, Estellé, et al. 2012)), segmental duplication (Cheng et al. 2005; Alkan et al. 2009) and tandem repeats filters (http://genome.ucsc.edu), as well as the requirement of orthology to chimp (suppl. fig. S13).

The analysis was performed on 3kb sliding (step 1.5 kb) genomic windows. Windows were defined in physical distance since the presence of LTBS may affect the population-based estimates of recombination rate. In addition, we chose to define windows based on length rather than number of ISs because the density of SNPs is part of *NCD2*’s signature, and fixing the number of ISs may result in particularly short windows in regions under balancing selection, which is undesirable for *NCD* analyses (see Results). We excluded 3 kb windows with less than 10 ISs in any population (suppl. information S2, suppl. figs S11 and S17) or less than 500 bp with orthology in chimp (1.6%); the two criteria combined resulted in the exclusion of 2.2% of scanned windows. Finally, to show that undetected short duplications do not contribute substantially to our candidates we analyzed an alternative modern human genome-wide dataset, sequenced to an average coverage of 20x-30x per individual (Meyer et al. 2012; Prüfer et al. 2013). After all these filters, we analyzed the 1,657,989 windows that remained in all populations, covering 2,145,937,383 base pairs (suppl. fig. S13).

#### Defining significant and outlier windows

We defined two sets of windows with signatures of LTBS, which collectively we consider our *candidate windows*. *Significant windows* are defined as those for which the observed *NCD2* value is lower than all values obtained from 10,000 neutral simulations with the same number of ISs. The matching by number of ISs aims to account for the higher variance in the statistic (in any SFS statistic) in windows with low numbers of informative sites (suppl. figs. S11 and S18). Thus, all significant windows have the same *p*-value (*p* < 0.0001).

In order to quantify how far the *NCD2* value of each window is from neutral expectations, we quantified the number of standard deviations a window’s *NCD2* value lies from the mean of the simulated distribution under neutrality (see suppl. information S2). For that, we defined a standardized distance measure between the observed *NCD2* (for a queried window) and the mean of the *NCD2*values for the 10,000 simulations with the matching number of ISs:

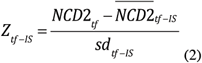

,where *Z*_*tf-IS*_ is the standardized *NCD2*, conditional on the value of IS*, NCD2*_*tf*_ is the *NCD2* value with a given *tf* for the *n-th* empirical window,*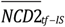* is the mean *NCD2* for 10,000 neutral simulations for the corresponding value of IS, and *sd*_*tf-IS*_ is the standard deviation for 10,000 *NCD2* values from simulations with matching ISs. *Z*_*tf-IS*_ allows the ranking of windows for a given *tf*, while taking into account the residual effect of IS number on *NCD2*_*tf*_, as well as a comparison between the rankings of a window considering different *tf* values. *Outlier windows* are defined based on an empirical *p*-value that is attributed to each window as a function of *Z*_*tf-IS*_ values for each *tf*. The empirical p-value is calculated by ranking the windows in terms of *Z*_*tf-IS*_ values and dividing the ranking position by the number of scanned window (i.e, the lowest empirical p-value is 1/1,657,989 = 5.97.10^−7^). Windows with empirical *p*-value ≤ 0.0005 (i.e., the 829 most extreme windows) were defined as *outlier windows.* With a few exceptions, outlier windows are a subset of significant windows. The coordinates for these windows are provided in suppl. table S5. For simplicity, all candidate windows that were identified for multiple *tf* values had an *assigned tf value*, defined as the one that minimizes the empirical *p*-value for a given window (suppl. information S2).

### Properties of candidate windows

#### Archaic introgression and ectopic gene conversion

We evaluated two potentially confounding biological factors. To assess ectopic gene conversion, we tested whether candidate genes have elevated number of paralogs in the same chromosome. To assess archaic introgression we calculated the proportion of European SNPs in candidate windows that are potentially of archaic (Neandertal or Denisova) origin. Details are provided in suppl. information S3.

#### Gene Ontology and tissue-specific expression enrichment analyses

We analyzed protein-coding genes overlapped by one or more candidate windows. GO and tissue of expression enrichment analyses were performed using GOWINDA (Kofler & Schlötterer 2012), which corrects for gene length-related biases and/or gene clustering. GO accession terms were downloaded from the GO Consortium website. We ran analyses in *mode:gene* (which assumes that all SNPs in a gene are completely linked) and performed 100,000 simulations for FDR (false discovery rate) estimation (significance at FDR ≤ 0.05). A minimum of three genes in the enriched category was required. For tissue-specific expression analysis we used Illumina BodyMap 2.0 (Derrien, Johnson, et al. 2012) expression data for 16 tissues, and considered genes significantly highly expressed in a particular tissue when compared to the remaining 15 tissues using the DESeq package (Anders & Huber 2010), as done in Sankararaman et al. (2014). Details in suppl. information S4.

#### SNP annotations

Functional annotations for SNPs were obtained from ENSEMBL-based annotations on the 1000G data and SNPs were categorized SNPs as intergenic, genic, exonic, regulatory, synonymous, and non-synonymous. For each candidate window, we sum the number of SNPs in each category, and then across candidate windows. To compare with non-candidate windows, we performed 1,000 re-samplings of the number of candidate windows from the set of all background (i.e. scanned) windows. Empirical *p*-values were obtained by comparing proportions of SNPs in each category to those obtained in 1,000 re-samplings (Figure 5 and suppl. table S5). Because we considered the sum across windows, and counted each SNP only once, results should be insensitive to window length (as candidate windows were merged). Details in suppl. information S4.

#### Splicing variants, monoallelic expression (MAE), immune-related genes

To test whether candidate genes have a larger number of transcripts than controls, while controlling for gene length, we: 1) divided candidate genes into quantiles of gene length; 2) sampled a set of genes (matching candidate genes in quantile bin; 3) calculated the mean and median number of transcripts for each set; 4) repeated this process 1,000 times and calculated the empirical one-tailed *p*-value. The number of transcripts per gene was obtained from Ensembl Biomart. For this analysis, to reduce complexity we used the set of candidate genes shared by populations within a continent (265 or 1594 for outlier and significant sets, respectively).

To test for enrichment for genes with mono-allelic expression (MAE), we quantified the proportions of candidate genes with MAE or bi-allelic expression in Savova et al. (2016) and compared these proportions to those in all genes analyzed (one-tailed Fisher’s test.) The same procedure was adopted to test for enrichment of immune-related genes among our sets, using a list of 386 keywords from the Comprehensive List of Immune Related Genes (https://immport.niaid.nih.gov/immportWeb/queryref/immportgene/immportGeneList.do).

Gene Cards was used to obtain basic functional information about genes. The GWAS catalog (Welter et al. 2014) was used to search for genome-wide phenotype associations included in the discussion (we only report “strong associations", i.e, when there is at least one SNP with *p* < 10-8). All statistical analyses and figures were performed in R (Development Core Team 2009) (scripts available on https://github.com/bbitarello/NCV_dir_package and NCD code available in https://github.com/bbitarello/NCD-Statistics).

## Results

### Power of NCD2 to detect LTBS

We evaluated the sensitivity and specificity of *NCD* by benchmarking its performance using simulations under demographic scenarios inferred for African, European, and Asian human populations and under a neutral or an overdominant model (see Methods). We explored the influence of parameters that can affect the power of *NCD*: time since onset of LTBS (*Tbs*), frequency equilibrium defined by selection coefficients (*f*_eq_), demographic history of the sampled population, *tf* used in *NCD* calculation and length of the genomic region analyzed (*L*). For simplicity, we averaged power estimates across African and European demographic models (Asian populations were not considered due to low power, see suppl. information S1), *L* and *Tbs*; these averages reflect general changes in power driven by individual parameters. For simplicity, we initially discuss cases where *tf* = *f*_eq_, and later relax this condition. We focus on *NCD2* (Table 1) power results and discuss some key points below. The complete set of power results, including *NCD1*, is presented in suppl. table S1.

**Table 1.** Power for simulations under the African and European demographic models. Power at false positive rate (FPR) = 5%. Simulations with *L* = 3 kb. *Tbs*, time in mya since onset of balancing selection; *f*_eq_, equilibrium frequency in the simulations. Power on additional conditions is presented on suppl. table S1.

|  |  | Africa |  |  |  |  |  | Europe |  |  |  |  |  |
| --- | --- | --- | --- | --- | --- | --- | --- | --- | --- | --- | --- | --- | --- |
| *Tbs* | $f_{eq}$ | *NCD2* | | | *NCD1* | | | *NCD2* | | | *NCD1* | | |
|  |  | *tf* |  |  | *tf* |  |  | *tf* |  |  | *tf* |  |  |
|  |  | 0.5 | 0.4 | 0.3 | 0.5 | 0.4 | 0.3 | 0.5 | 0.4 | 0.3 | 0.5 | 0.4 | 0.3 |
| 5 | 0.5 | 0.96 | 0.94 | 0.84 | 0.93 | 0.91 | 0.39 | 0.97 | 0.95 | 0.84 | 0.92 | 0.85 | 0.20 |
| 5 | 0.4 | 0.94 | 0.94 | 0.89 | 0.89 | 0.89 | 0.67 | 0.95 | 0.94 | 0.91 | 0.85 | 0.83 | 0.60 |
| 5 | 0.3 | 0.90 | 0.91 | 0.93 | 0.72 | 0.80 | 0.84 | 0.84 | 0.86 | 0.89 | 0.47 | 0.57 | 0.74 |
| 3 | 0.5 | 0.91 | 0.88 | 0.68 | 0.86 | 0.80 | 0.24 | 0.93 | 0.89 | 0.68 | 0.81 | 0.69 | 0.15 |
| 3 | 0.4 | 0.88 | 0.86 | 0.76 | 0.78 | 0.78 | 0.56 | 0.89 | 0.88 | 0.79 | 0.74 | 0.71 | 0.46 |
| 3 | 0.3 | 0.75 | 0.77 | 0.81 | 0.56 | 0.64 | 0.71 | 0.73 | 0.76 | 0.80 | 0.39 | 0.48 | 0.63 |

#### Time since the onset of balancing selection (*Tbs*) and sequence length

Signatures of LTBS are expected to be stronger for longer *Tbs*, because time to the most recent common ancestor is older and there will have been more time for linked mutations to accumulate and reach intermediate frequencies. Accordingly, power to detect LTBS is high with *Tbs* = 3 (0.91) and 5 (0.96) and low for 1 mya (0.42, averaged across populations, *L* and *f*_eq_) (suppl. figs. S3-S8, suppl. table S1). Further, in the absence of epistasis the long-term effects of recombination result in narrow signatures with old *Tbs*. Indeed, our simulations show that narrower regions yield highest power ( on average 13.6% higher for 3 than for 12kb, when using *NCD2(0.5)*, *Tbs* = 5 mya; suppl. figs. S3-S8, suppl. table S1). Thus, we focus the remaining power discussion on old balancing selection (3 or 5 mya) in regions of 3kb.

#### Demography and sample size

Power is similar for samples simulated under African and European demographic histories for *NCD2* (table 1) but considerably lower under the Asian one, possibly due to demography (suppl. table S1, suppl. figs. S3-S8). While power estimates may be influenced by the particular demographic model used, we nevertheless focus on African and European populations, which by showing similar power allow fair comparisons between them. Although the power values discussed here are for samples of 100 chromosomes, *NCD2* performs well even for sample sizes as small as 20 (suppl. table S1 and suppl. information S1).

#### Simulated and target frequencies

So far, we have only discussed cases where *tf* = *f*_eq_, which should favor the performance of *NCD*. Accordingly, under this condition *NCD* has very high power: 0.96, 0.94, and 0.89 for *f*_eq_ = 0.5, 0.4, and 0.3, respectively (averaged across *Tbs* and populations, 3kb, Table 1). However, in practice there is no prior knowledge about the *f*_eq_ of balanced polymorphisms, so we also evaluate the power of *NCD* when *tf* differs from the *f*_eq_. When *f*_eq_ = 0.5, average power is high for *tf* = 0.5 and 0.4 (≥ 0.94), but lower for *tf* = 0.3 (0.83, Table 1). Similar patterns are observed for other simulated *f*_eq_ (Table 1). Therefore, *NCD2* is overall well-powered both when *f*_eq_ is the same as *tf* and also when *f*_eq_ ≠ *tf*. In any case, the closer *tf* is to *f*_eq_, the higher the power, so when possible it is desirable to perform tests across a range of *tf.*

#### *NCD2* in comparison to other methods

We compared *NCD2* to two statistics commonly used to detect balancing selection (Taj*D* and HKA), and two recently proposed high-powered statistics: the composite likelihood-based measure *T*_2_ (DeGiorgio et al. 2014) and ß (Siewert & Voight 2017). The *T*_2_ statistic, similarly to *NCD2*, considers both the SFS and FDs, whereas ß only considers the SFS. When *f*_eq_ = 0.5, *NCD2(0.5)* has the highest power: for example, for the African demographic scenario (using *Tbs* = 5 mya, and 3 kb windows) *NCD2(0.5)* has a power of 0.96 (the highest among other tests is 0.94, for *T*_2_) but the difference in power is highest when *f*_eq_ departs from 0.5. For *f*_eq_ = 0.4, *NCD2(0.4)* power is 0.93 (compared to 0.90 for Taj*D* and *T*_2_, and lower for the other tests). For *f*_eq_ = 0.3, *NCD2(0.3)* power is 0.93 (compared to 0.90 for ß, 0.89 for *T*_2_ and lower for the other tests). These patterns are consistent in the African and European simulations (Figure 2, suppl. fig. S10), where *NCD*2 has greater or comparable power than other available methods.

*NCD2* is simple to implement and fast to run, and performs slightly or substantially better than all other methods tested (Figure 2, suppl. fig. S10). Altogether, the advantage of *NCD2* over classic neutrality tests and ß is its higher power, especially when *f*_eq_ departs from 0.5; the main advantage over *T*_2_ is simplicity and speed of implementation. We note that *NCD2* can be computed for particular loci, even in the absence of genome-wide data (which, if present, allows an empirical approach to define significance) or a demographic model (which, if present, allows simulations to define significance). Also, *NCD2* outperforms *NCD1* (see suppl. information S1), illustrating the gain in power by incorporating FDs in the *NCD* statistic, which is also more powerful than combining *NCD1* and HKA (suppl. table S1 and information S1). Because *NCD2* outperforms *NCD1*, we used it for our scan of human populations. However, *NCD1* also outperforms several of the existing methods (suppl. table S1) and is a good choice when outgroup data is lacking.

### Identifying signatures of LTBS in human genomes

The ideal window size will vary depending on species demographic properties and sampling. Informed by our power analyses, we computed the highly powered *NCD2* in sliding windows of 3 kb. These were defined as sliding windows in a genome-wide framework, to ensure overlap with the narrow signatures of balancing selection. We note though that alternative approaches to define the windows, such as windows centered on each IS, are also possible. After filtering (see Methods), we analyzed about 81% of the autosomal genome (suppl. fig. S13), overlapping 18,633 protein-coding genes. Between 6,226 and 6,854 (0.37-0.41%) of the scanned windows in each population have a lower *NCD2(0.5)* value than any of the 10,000 neutral simulations (*p* < 0.0001), placing them in the *significant* category (table 2). The proportions are similar for *NCD2(0.4)* and *NCD2(0.3)* (table 2). In each population, the union of significant windows considering all *tf* values spans, on average, 0.6% of the windows (table 2) and 0.77% of the base pairs.

**Table 2.**
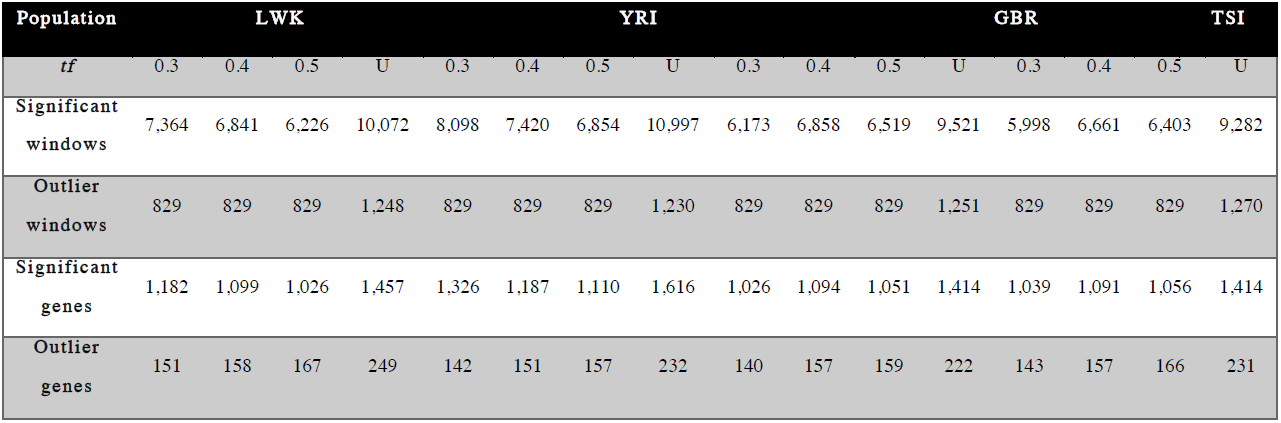
Candidate windows and protein-coding genes across populations. Note. – Significant and outlier genes and windows, see main text. U, union of windows considering three *tf* values. Total number of queried windows per population is 1,657,989. Union of all candidate genes is 2,348 (significant) and 402 (outlier).

| Population | LWK |  |  |  | YRI |  |  |  | GBR |  |  |  | TSI |  |  |  |
| --- | --- | --- | --- | --- | --- | --- | --- | --- | --- | --- | --- | --- | --- | --- | --- | --- |
| *tf* | 0.3 | 0.4 | 0.5 | U | 0.3 | 0.4 | 0.5 | U | 0.3 | 0.4 | 0.5 | U | 0.3 | 0.4 | 0.5 | U |
| **Significant windows** | 7,364 | 6,841 | 6,226 | 10,072 | 8,098 | 7,420 | 6,854 | 10,997 | 6,173 | 6,858 | 6,519 | 9,521 | 5,998 | 6,661 | 6,403 | 9,282 |
| **Outlier windows** | 829 | 829 | 829 | 1,248 | 829 | 829 | 829 | 1,230 | 829 | 829 | 829 | 1,251 | 829 | 829 | 829 | 1,270 |
| **Significant genes** | 1,182 | 1,099 | 1,026 | 1,457 | 1,326 | 1,187 | 1,110 | 1,616 | 1,026 | 1,094 | 1,051 | 1,414 | 1,039 | 1,091 | 1,056 | 1,414 |
| **Outlier genes** | 151 | 158 | 167 | 249 | 142 | 151 | 157 | 232 | 140 | 157 | 159 | 222 | 143 | 157 | 166 | 231 |

Next, we defined as *outlier* windows for each *tf* those with the most extreme signatures of LTBS (in the 0.05% lower tail of the respective *Ztf-IS* distribution, see Methods). This more conservative set contains 829 outlier windows for each population and *tf* value (table 2), which cover only ∼ 0.09% of the base pairs analyzed and are largely included in the set of significant windows. Significant and outlier windows are collectively referred to as *candidate windows*.

#### Reliability of candidate windows

Significant windows are enriched both for polymorphic sites (Figure 3A-B) and intermediate-frequency alleles, and the SFS shape reflects the *tf* for which they are significant (Figure 3C-D). Although expected, because these were the patterns used to identify these windows, this shows that significant windows are unusual in both signatures. Also, as expected with LTBS, the significant windows are largely shared across populations (see below). The striking differences of significant windows with respect to the background distribution, combined with the fact that neutral simulations do not have *NCD2* values as low as those of the significant windows, precludes relaxation of selective constraint as a an alternative explanation to their signatures (Andrés et al. 2009). Relaxation of purifying selection can shift slightly the SFS (mostly for the functional sites, e.g. non-synonymous), but it cannot explain our significant windows because their SFS is strikingly different from the genome-wide, background distribution (which represents mostly neutral regions) and from simulations run under neutrality.

**Fig. 3.**
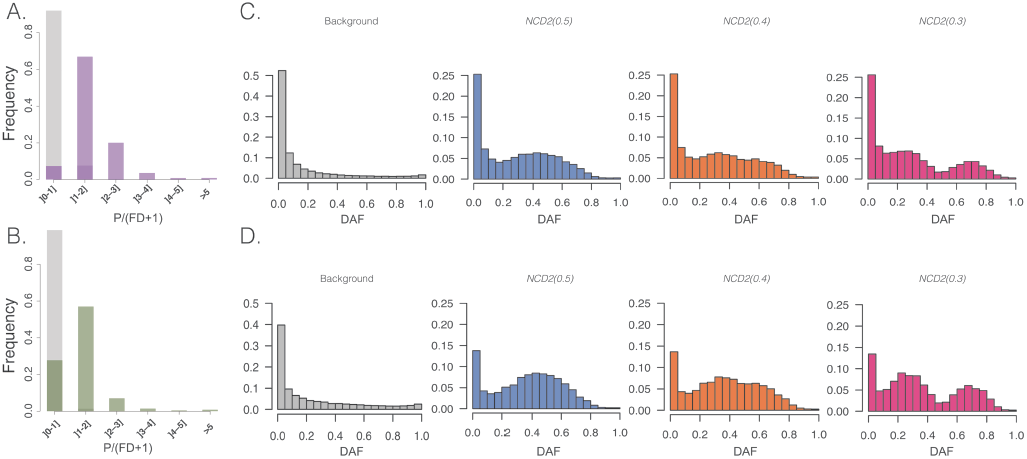
Polymorphism-to-divergence and SFS. **(A, B)** SNPs/(FDs+1) for LWK **(A)** and GBR **(B)** populations. SNPs/(FDs+1) measures the proportion of polymorphic-to-divergent sites for the union of significant windows for all *tf* (purple, green) compared to all scanned windows (gray). **(C-D)** SFS in LWK **(C)** and GBR **(D)** of all scanned windows in chr1 (gray), significant windows for *NCD2(0.5)* (blue), *NCD2(0.4)* (orange), *NCD2(0.3)* (pink). DAF, derived allele frequency.

Windows with low *NCD2* values have similar coverage to the rest of the genome, i.e, they are not enriched in unannotated, polymorphic duplications. Also, candidate windows are not only not enriched in high-coverage regions: they show a depletion of them (suppl. fig. S14) (Mann-Whitney U two-tail test; *p* < 0.02 for *tf* = 0.5 and *tf* = 0.4 for GBR and TSI). We also examined whether these signatures could be driven by two biological mechanisms other than LTBS: archaic introgression into modern humans and ectopic gene conversion (among paralogs). These mechanisms can increase the number of polymorphic sites and (in some cases) shift the SFS towards intermediate frequency alleles. We find introgression is an unlikely confounding mechanism, since candidate windows are depleted in SNPs introgressed from Neandertals (suppl. fig. S16, suppl. table S4, suppl. information S3). Also, genes overlapped by significant windows are not predicted to be affected by ectopic gene conversion with neighboring paralogs to an unusually high degree, with the exception of olfactory receptor genes (suppl. fig. S15, suppl. information S3).

The candidate windows are enriched in heterozygous positions in the genomes of archaic humans (Neandertal and Denisova, all p-values < 0.001). This shows that, as expected, the polymorphisms in identified candidates tend to be older than those in neutral regions of the genome, predating the split between humans and archaic hominins (suppl. information S6, suppl. fig. S22).Thus, candidate windows represent a catalog of strong candidate targets of LTBS in human populations.

#### Assigned *tf* values

Many windows were significant for more than one *tf*. For these cases, we used the Ztf-IS statistic (Equation 2) to identify which *tf* provides the strongest support for LTBS (i.e., for which *tf* the departure from neutral expectations was greatest). In this way, we could assign a *tf* to each significant window. On average ∼53% of the candidate windows are assigned to *tf* = 0.3, 27% to *tf* = 0.4 and 20% to *tf* = 0.5 (suppl. table S3 and information S2).

#### Non-random distribution across chromosomes

Candidate windows are not randomly distributed across the genome. Chromosome 6 is the most enriched for signatures of LTBS, contributing, for example, 10.2% of significant and 25% of outlier windows genome-wide for LWK while having only 6.4% of analyzed windows (suppl. fig. S12). This is explained by the MHC region (Figure 4A), rich in genes with well-supported evidence for LTBS. Specifically, 10 HLA genes are among the strongest candidates for balancing selection in all four populations, most of which have prior evidence of balancing selection (suppl. table S4, suppl. information S4 and S5).

**Fig. 4.**
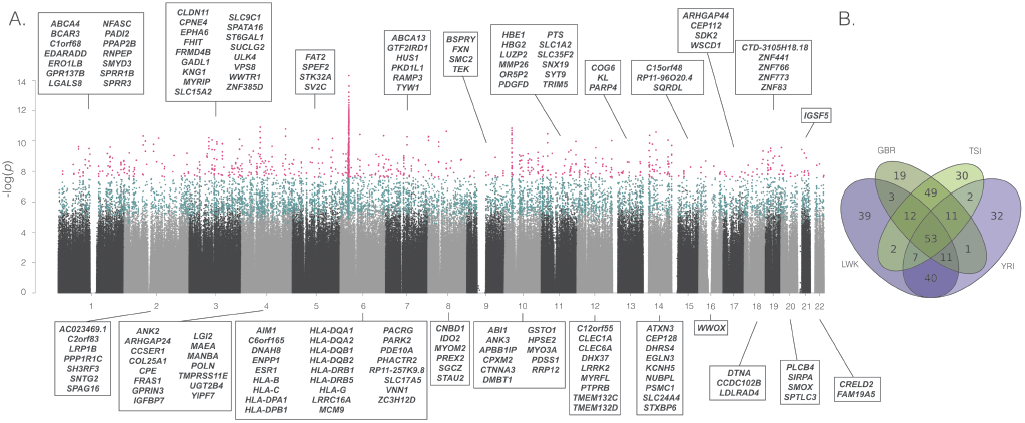
Manhattan plot and population sharing. **(A)** Manhattan plot of all scanned windows, for one analysis (*NCD2(0.5)* for LWK). *y*-axis, *p*-value (log-scale) based on *Ztf-IS*. *x*-axis, ordered location of analyzed windows on the genome. Each point is a scanned (grey and black), significant (blue) or outlier (pink) window. Names of outlier protein-coding genes are provided, sorted by name. Significant windows were defined based of simulations, not on *Ztf-IS*. (*Ztf-IS* is used to rank even those with *p* < 0.0001) **(B)** Venn diagram showing the overlap in signatures of the 167 outlier genes annotated in **(A)** with other populations.

### Biological pathways influenced by LTBS

To gain insight on the biological pathways influenced by LTBS, we focused on protein-coding genes containing at least one candidate window (222-249 outlier and 1,404-1,616 significant genes per population), and investigated their annotations (all results are presented in suppl. table S2). Regarding functional categories, significant genes are overrepresented in 28 GO categories, 24 of which are shared by at least two populations and 18 by four populations (suppl. table S2). Thirteen categories are immune-related according to a list of 386 immune-related keywords from ImmPort (Methods). The more stringent sets of outlier genes are enriched for 28 GO categories (21 shared by all four populations), 18 of which are immune-related. Furthermore, in both sets several of the remaining enriched categories are directly related to antigen presentation although not classified as immune-related (e.g. “ER to golgi transport vesicle membrane", “integral to membrane”). Among the non immune-related categories are “sarcolemma", “epidermis development", “keratin filament” and “negative regulation of blood coagulation".

When classical HLA genes are removed from the analyses (suppl. information S4), only two categories remain enriched: “sarcolemma” (in YRI) and “epidermis development” (GBR), but the small set of genes per population hampers power. For the significant windows, “antigen processing and presentation of endogenous peptide antigen via MHC class I” remains significantly enriched (driven by *TAP1*, *TAP2, ERAP1* and *ERAP2*; suppl. table S2). Significant windows remain enriched in categories related to the extracellular space – “extracellular regions", “integral to membrane” – as in other studies (Andrés et al. 2009; DeGiorgio et al. 2014; Key, Teixeira, et al. 2014) – and “keratin filament". These categories are not immune-related *per se*, but they represent physical barriers to the invasion by pathogens. This indicates that LTBS maintains advantageous diversity in numerous defense-related genes other than classical HLA genes.

Overall, 33% of the outlier (and 31% of the significant) genes have at least one immune-related associated GO category, compared to only 24% of scanned genes (see Methods). These results collectively suggest that immunity and defense are frequent targets of LTBS, although a large fraction of the candidates for LTBS have non-immune related functions or indirect connections with immunity hitherto unknown.

### Functional annotation of SNPs in candidate windows

We next tested whether LTBS preferentially targets SNPs at particular types of functional sites. To do so we investigated the overlap of candidate windows with different classes of functional annotations in the human genome (Methods and suppl. information S4), and tested the hypothesis of enrichment of certain classes of sites within our sets of candidate windows, when compared to sets of randomly sampled windows from the genome.

SNPs in outlier windows are disproportionally represented in protein-coding exons in all the populations (*p* ≤ 0.001, one-tail test; Figure 5 and suppl. table S5). The protein-coding enrichment is even stronger when considering only SNPs within genes, which both in outlier (*p* < 0.001) and significant windows (*p* ≤ 0.003) are strongly enriched in protein-coding exons (Figure 5). Within the protein-coding exons, outlier windows in Africa (*p* ≤ 0.022) and significant windows in all populations (*p* ≤ 0.037) are enriched for non-synonymous SNPs (Figure 5). These observations show that our candidate targets of LTBS tend to be enriched in exonic and potentially functional (amino-acid altering) SNPs.

**Fig. 5.**
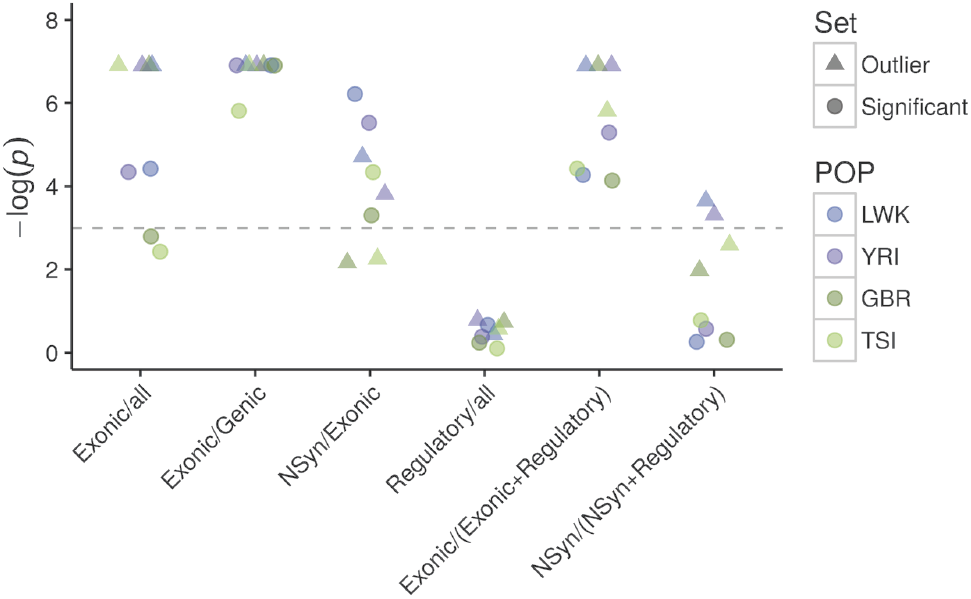
Enrichment of classes of sites amongst candidate windows. Dashed lines mark the *p* = 0.975 (bottom) and *p* = 0.025 (top) thresholds for the one-tailed *p*-values (hypothesis: enrichment). NSyn, nonsynonymous; all, Genic + Intergenic + Regulatory. The annotation is based on Ensembl variant predictor (suppl. information S3). *p* < 0.001 was treated as 0.001 to avoid infinite values.

Conversely, outlier and significant windows have no excess of SNPs annotated as regulatory (*p* ≥ 0.458 in all populations, Figure 5). When we explicitly compared protein-coding exons vs. regulatory sites by restricting our analysis to sites in these two categories, outlier windows have an excess of exonic SNPs (*p* ≤ 0.003). The same is true for significant windows (*p* ≤ 0.016; Figure 5). When only nonsynonymous and regulatory sites are considered, we see enrichment for LWK and YRI for the outlier windows (*p* ≤ 0.036, Figure 5) but not for the significant windows (*p* ≥ 0.458 for all populations, Figure 5), although the two analyses that consider nonsynonymous SNPs are likely underpowered due to low SNP counts (suppl. table S5).

Finally, results using more detailed RegulomeDB annotations generally agree with the observation of lack of enrichment of regulatory sites in our candidate windows (*p* ≥ 0.121 for a one tail test for enrichment for RegulomeDB1+3 for SNPs with MAF ≥ 0.2) (suppl. table S5). Although perhaps limited by the quality of the annotation of regulatory sites and the low power associated to small SNP counts for nonsynonymous variants, we do not have strong evidence that LTBS in human populations has preferentially shaped variation at sites with a role in gene expression regulation.

### Overlap across populations

On average 86% of outlier windows in a given population are shared with another population (79% for significant windows), and 77% with another population within the same continent (66% for significant ones) (suppl. fig. S18). The sharing is similar when *tf* are considered separately (suppl. figs. S19 and S20). Therefore, there is also considerable overlap of candidate protein-coding genes across populations: e.g. in LWK (*tf* = 0.5), 76.6% of outlier genes are shared with at least another population, and 66% are shared with YRI (89% and 77% for significant genes; Figure 4B). In fact, on average 44% of outlier genes for a given population are shared across *all* populations and 78.7% are shared by a same-continent population (50% and 77% for significant genes; suppl. fig. S21).

#### Candidate genes in more than one population

Instances where signatures of LTBS are not shared between populations may result from changes in selective pressure, which may be important during fast, local adaptation (de Filippo et al. 2016). On the other hand, loci with signatures of LTBS across human populations are more likely to represent stable selection. We considered as “shared” those candidate protein-coding genes (from the union of candidate windows for all *tf*) that are shared by all populations (suppl. table S4). We considered as “African” those shared between YRI and LWK (but neither or only one European population), and “European” those shared between GBR and TSI (but neither or only one African population). We note that these designations do not imply that genes referred to as “African” or “European” are putative targets of LTBS for only one continent (partially because there are some power differences between Africa and Europe, table 1). The 79 African, 84 European and 102 shared outlier genes add up to 265 genes in total (∼1.4% of all queried genes) and the 458 African, 400 European, and 736 shared significant genes add up to 1,594 (∼8.5% of all queried genes; suppl. table S4). Several of them have been detected in other studies, but the vast majority are novel candidates (suppl. information S4 and S5).

### Expression of candidate genes

Candidate genes are disproportionally expressed in a number of tissues: lung, adipose tissue, adrenal tissue, kidney, and prostate (FDR ≤ 0.05, suppl. table S2). Also, candidate genes have an unusually large number of different transcripts: outlier genes shared by at least two populations have on average 8.69 transcripts (compared to 7 in the controls after controlling for length), with a similar pattern holding for significant genes (8.73 compared to 7.1 in background control genes).These are both significantly higher than controls even after controlling for gene length (*p* < 0.001 in both cases for a one-tail test).

Genes with mono-allelic expression (MAE) – i.e, the random and mitotically stable choice of an active allele for a given locus – were enriched among the small set of genes previously reported to be under LTBS (Savova et al. 2016). Our observations are in agreement with these findings, with 64% and 62% of the outlier and significant genes shared by at least two populations having MAE status (Savova et al. 2016), compared to only 41% for genes without signatures of LTBS (*p <* 1.12-6 Fisher Exact Test, one-sided).

## Discussion

### Limitations of NCD statistics

LTBS produces signatures that are unexpected under neutrality. Still, all tests for balancing selection are affected by demographic history, in particular strong population substructure, which can result in an excess of polymorphism that, in certain cases (e.g. with similar contribution of the two subpopulations, or with positive selection) may be at intermediate frequencies. We benefit from the extensive prior work on the 1,000 Genomes populations showing absence of substructure and providing a parameter-rich demographic model, which can be used for hypothesis testing. Some types of migration and introgression can also result in genome-wide or local excess of diversity and, under positive selection, also intermediate frequencies. We show that Neandertal introgression has a minimal effect on our inferences of LTBS. However, as with any neutrality test, applying *NCD* to other species requires consideration of the demographic history of the population to weight the possibility of alternative explanations. If the demographic history is unknown, a purely empirical approach allows the identification of putative targets of balancing selection. Nevertheless, substructure, migration and introgression should to be investigated as alternative explanations, using additional methods (e.g. LD-based).

Certain selective regimes can also generate signatures that are similar to those of LTBS. For example, incomplete or soft sweeps can produce an excess of intermediate frequency variants (Hermisson & Pennings 2017) that could be mistaken as evidence for balancing selection when only polymorphisms are used. Still, because none of these types of selection increase the time to the most recent common ancestor and density of polymorphisms, they do not confound *NCD2*. Recent balancing selection, on the other hand, is difficult to distinguish from those two selective scenarios, so we focused our analyses and interpretations on LTBS. Of note, by definition *NCD* requires pre-defining a *tf*, which in practice is unknown, but this can be easily addressed by running the test with different *tf* (whose results correlate strongly) and combining results. So, overall, and despite these limitations, *NCD* is a powerful, simple and fast method to identify the signatures of LTBS in polymorphism data, which we expect will be used in model and non-model organisms.

### The targets of LTSB in the human genome

Using simulation-based and outlier approaches, we uncovered windows with signatures of LTBS in humans. We showed that these windows are unlikely to be affected by technical artifacts or biological processes other than LTBS, such as introgression from archaic hominins. On average, across populations, 0.6% of the windows in a population meet our criterion of significance based on neutral simulations. These windows contain on average 0.77% of the base-pairs and 1.6% of the SNPs in the analyzed genome per population, and although they amount to a low proportion of the genome, on average 7.9% of the protein-coding genes contain at least one significant window (considering UTRs, introns and protein coding exons). For the outlier windows (those in the 0.05% lower tail of the *NCD2* genomic distribution), on average 1.2% of genes show some evidence of selection. These proportions are similar when requiring that a significant or outlier window be shared by at least two same-continent populations (8.5% and 1.4%, respectively). We note that although these sets probably include some false positives, our method is only sensitive to very old balancing selection, so in practice we are likely underestimating the total influence of balancing selection in human genomes. For instance, a more widespread influence of younger instances of balancing selection, which cannot be detected by our methods, could in principle be detected by methods able to distinguish the signatures of recent balancing selection from those of incomplete sweeps or selection on standing variation. Alternatively, recent balancing selection targeting particular pathways or gene networks could in principle be detected by gene set enrichment analyses as described in Daub et al. (2013) in the context of polygenic adaptation.

We identified a number of previously known targets of LTBS, but also many new ones: almost 70% of the outlier genes shared at least by another same-continent population (and 90% of the significant ones) are novel. Many of the candidate genes show strongest evidence for LTBS at *tf* values different from 0.5. This is expected, for instance, under asymmetric overdominance, and highlights the importance of considering selective regimes with different frequencies of the balanced polymorphism.

### Functional properties of SNPs in candidate windows

In this study, we confirm cases where protein-coding regions are the likely targets of selection, such as *HLA-B* and *HLA-C* (Hughes & Nei 1988), as well as cases where regulatory regions are probably targeted, such as *UGT2B4, TRIM5* (Cagliani et al. 2010; Sun et al. 2011). While LTBS has been proposed to play an important role in maintaining genetic diversity that affects gene expression (Leffler et al. 2013; Savova et al. 2016), we find that regulatory SNPs are underrepresented within the candidate regions. This does not imply that there are no regulatory SNPs under LTBS, but rather that with existing annotations (which are less precise for regulatory than protein-coding sites) they are not enriched within candidate windows. Overall, we found an enrichment of exonic and of non-synonymous SNPs in the candidate windows, suggesting an overrepresentation of potentially functional SNPs within selected regions. This is compatible with two scenarios: (a) direct selection on protein-coding sites or (b) accumulation of functional (including slightly deleterious) variants as a bi-product of LTBS.

### Overlap with previous studies

Whereas the various scans for positive selection show a remarkably low overlap with respect to the genes they identify, 34% of our outlier genes (11% of significant ones) had evidence of LTBS in at least one previous study (Andrés et al. 2009; Leffler et al. 2013; DeGiorgio et al. 2014). Remarkably, 47% of the shared outliers across all four populations (17% of the shared significant ones) have been detected in at least one previous study, and the proportions are similar even when classical HLA genes are removed (39 and 16% overlap, respectively). This is a high degree of overlap, considering the differences in methods and datasets across studies.

### Properties of candidate genes

Below we discuss some of the candidate genes (suppl. table S4), highlighting the variety of biological functions and known genetic associations potentially shaped by LTBS in humans.

#### Gene expression

Candidate genes have on average a larger number of transcripts than other genes. As examples, *CLDN11*, *ROBO2*, *ESR1* or *PRKCE* (discussed below) have 10-15 different transcripts, a higher number than at least 75% of scanned genes. It is possible that these genes benefit from high levels of diversity, contributed both by genetic diversity and transcript diversity. Also in agreement with previous findings we find a significant excess of MAE genes among targets of LTBS. This excess is not driven by HLA genes and supports the claim for a biological link between MAE and balancing selection (Savova et al. 2016). Heterozygosity in a MAE gene could lead to cell-to-cell heterogeneity within same-cell clusters, which could be potentially advantageous (Savova et al. 2016), particularly in the case of cell-surface proteins. Some of these MAE genes found in our study, and not previously detected in scans for LTBS, are involved in immunity/defense barriers (e.g. *IL1RL1*-*IL18R1, FAM114A1, EDARADD, SIRPA, TAS2R14)*, oxygen transport and hypoxia (e.g. *PRKCE*, *HBE1, HBG2*, *EGLN3),* or reproduction (e.g. *CLDN11).*

#### Oxygen transport and response to hypoxia

Among the outlier genes with MAE we find *EGLN3,* which is significantly upregulated under hypoxia (Escribese et al. 2012) and plays a role in skeletal muscle differentiation (Fu et al. 2007). The encoded protein hydroxylates the product of *EPAS1*, a gene shown to harbor variants responsible for human adaptation to high altitude in Tibet (Yi et al. 2013). We also find members of the ß-globin cluster (*HBE1*-*HBG2*) that are involved in oxygen transport and have strong associations to hemoglobin levels and ß-thalassemia (Danjou et al. 2015), and for which there is also evidence for recent positive selection in Andean (*HBE1*, *HBG2*) or Tibetan (*HBG2*) populations (Bigham et al. 2010; Rottgardt et al. 2010; Yi et al. 2010). It is plausible that these genes have been under LTBS and undergone a shift in selective pressures in high-altitude populations, but further analyses are required to confirm this possibility. Another outlier gene, *PRKCE,* is also associated to hemoglobin levels and red blood cell traits (Astle et al. 2016).

#### Immunological function and defense barriers

It has long been argued that genes of immune function are prime candidates for LTBS. As expected, we detect several classical HLA with known signatures of LTBS. However, many non-HLA candidates from our set of outlier genes have immunological functions. We confirm signatures of LTBS in the *ABO* locus (suppl. information S5), a well-known case of LTBS in humans (Ségurel et al. 2012), and *TRIM5*, a gene with antiviral function (Cagliani et al. 2010). Among novel candidates, we find several genes involved in auto-immune disease. For example, *IL1RL1*-*IL18R1* have strong associations atopic dermatitis, an auto-immune disease (Hirota et al. 2012). *HLA-DQB2* mediates superantigen activation of T cells and is associated both to infectious (hepatitis B) and autoimmune diseases (e.g. Lee et al. 2012; Jiang et al. 2015). Two other significant genes for which there is prior evidence for LTBS, *ERAP1* and *ERAP2* (Andrés et al. 2010; Cagliani, Riva, et al. 2010), are associated with ankylosing spondylitis and psoriasis (e.g. Strange et al. 2010; Evans et al. 2011; Robinson et al. 2015). Finally, there are several associations to autoimmune disease and susceptibility to infections in the classical HLA genes that we identify. In brief, our results are consistent with the hypothesis that auto-immune disease is linked to natural selection favoring effective immune response against pathogens (Corona et al. 2010; Sironi & Clerici 2010).

Another important aspect of defense is the avoidance of poisonous substances. As suggested previously by studies on polymorphism in PTC receptors (Wooding et al. 2004), avoidance of bitterness might have been adaptive throughout human evolutionary history because several potentially harmful substances are bitter. The *TAS2R14* gene encodes for a bitter taste receptor, and in humans it has strong associations to taste perception of quinine and caffeine (Ledda et al. 2014), is considered a promiscuous receptor (Meyerhof et al. 2010; Karaman et al. 2016), and is one of the few bitter taste receptors that binds a vast array of compounds, and for which no common structure has been found (Behrens et al. 2004; Meyerhof et al. 2010). This entails diversity in the antigen binding portions of the receptors, which may be enhanced by balancing selection. Indeed, an elevated *dN/dS* ratio was reported for a cluster of bitter taste receptors which includes *TAS2R14* (Kosiol et al. 2008). To our knowledge, our study is the first in detecting signatures of LTBS in this gene.

#### Cognition

Several candidate genes are involved in cognitive abilities, or their variation is associated with diversity in related phenotypes. *KL* (life extension factor klotho) is a gene that has been associated to human longevity (Arking et al. 2002) and for which signatures of LTBS have been previously reported (DeGiorgio et al. 2014). In mice, decreased levels of klotho shorten lifespan (Welberg 2014). In humans, heterozygotes for the KL-VS variant show higher levels of serum klotho and enhanced cognition, independent of sex and age, than wild-type homozygotes. On the other hand, KL-VS homozygotes show decreased lifespan and reduced cognition (Dubal et al. 2014). If higher cognition is advantageous, overdominance for this phenotype can explain the signatures of balancing selection we observe, although the effect in lifespan itself could also influence the selective regime.

*PDGFD* encodes a growth factor that plays an essential role in wound healing and angiogenesis. PDGFD-induced signaling is crucial for human (but not mouse) proliferation of the neocortex due to neural stem-cell proliferation (Lui et al. 2014), a trait that underlies human cognition capacities. This gene has strong associations to coronary artery disease and myocardial infarction, which are related to aging.

Also among our outliers is a gene with a cognitive-related genetic association, *ROBO2,* which encodes a transmembrane receptor involved in axon guidance. Associations with vocabulary growth have been reported for variants in its vicinity (St Pourcain et al. 2014). *ROBO2* has signatures of ancient selective sweeps in modern humans after the split with Neandertals and Denisova (Peyrégne et al. 2017) on a portion of the gene (chr3:77027850-77034264) almost 40kb apart from the one for which we identified a signature of LTBS (chr3:76985072-76988072). The occurrence of both these signatures highlights the complex evolutionary history of this gene.

Associations of candidate genes with cognition are also exemplified by case-control and cohort studies linking polymorphisms in the estrogen receptor alpha (ER-α) gene, *ESR1*, to dementia and cognitive decline. Links between ER-α variants and anxiety and depression in women have been proposed but lack confirmation (reviewed in Sundermann et al. 2010). Interestingly, three other candidate genes (*PDLIM1,GRIP1, SMYD3)* interact with ER-α at the protein level (Szklarczyk et al. 2015), and two (*PDLIM1,GRIP1*) have strong association with suicide risk (Perlis et al. 2010; Mullins et al. 2014).

In genes like *KL*, where heterozygotes show higher cognitive abilities than homozygotes, cognition may be a driving selective force. This is a possible scenario in other genes, too. Still, given the complexity of brain development and function, it is also possible that cognitive effects of this variation are a byproduct of diversity maintained for other phenotypes. For example, MHC proteins and other immune effectors are believed to affect connectivity and function of the brain (reviewed in (Shatz 2009; Needleman & McAllister 2012), with certain alleles being clearly associated with autism disorder.

#### Reproduction

Among candidate genes, there is an enrichment for preferential expression in the prostate. There are also a number of outlier genes involved in the formation of the sperm. For example, *CLDN11* encodes a protein expressed in several tissues and crucial for spermatogenesis. Knockout mice for the murine homologue show neurological and reproductive impairment (Gow et al. 1999; Wu et al. 2012) and some variants in humans are strongly associated to prostate cancer.

*ESR1*, mentioned above, encodes an estrogen-activated transcription factor and leads to abnormal secondary sexual characteristics in females when defective (Quaynor et al. 2013). ER-α interacts directly with the product of *BRCA1* and has strong associations to breast cancer (Michailidou et al. 2013), breast size (Eriksson et al. 2012) and age of menarche. In males, it is involved in gonadal development and differentiation, and lack of estrogen and/or ER-α can lead to poor sperm viability (Lazari et al. 2009). Strikingly, this gene also has SNPs strongly associated to a diverse array of phenotypes, including height, bone mineral density (spine and hip), and sudden cardiac arrest (Rivadeneira et al. 2009; Aouizerat et al. 2011; Wood et al. 2014). Two other genes among our candidates are also part of the estrogen signaling pathway: *PLCB4* and *ADCY5* (which is strongly associated to birth weight). Estrogens are not only involved in reproductive functions (both in male and females), but also in several processes of neural (see above), muscular or immune nature, and the ER-α-estrogen complex can act directly on promoter regions of other genes. In this case, balancing selection could be explained by the high level of pleiotropy, including the function in male and female reproduction (if different alleles are beneficial in males than females).

### Conclusions

We present two new summary statistics, *NCD1* and *NCD2*, which are both simple and fast to implement on large datasets to identify genomic regions with signatures of LTBS. They have a high degree of sensitivity for different equilibrium frequencies of the balanced polymorphism and, unlike classical statistics such as Tajima’s *D* or the Mann-Whitney U (Andrés et al. 2009; Nielsen et al. 2009), allow an exploration of the most likely frequencies at which balancing selection maintains the polymorphisms. This property is shared with the ß and the likelihood-based *T*_2_ tests (DeGiorgio et al. 2014; Siewert & Voight 2017). We show that *NCD* is well-powered to detect LTBS within a complex demographic scenario, such as that of human populations. *NCD* can be applied to either single loci or the whole-genome, in species with or without detailed demographic information, and both in the presence and absence of an appropriate outgroup (*NCD2* and *NCD1*, respectively).

More than 85% of our outlier windows are shared across populations, raising the possibility that long-term selective pressures have been maintained after human populations colonized new areas of the globe. Still, about 15% of outlier windows show signatures exclusively in one sampled population, and 16 of these show opposing signatures (*OAS1, C15orf48, OR11A1, GSTO1, DHRS4, PABPC1, MYOZ3, OR6K5, KRTAP10-7, GRTP1, MINOS1, GANC, FBXO15, TIMM21, ZNF780A, MAPT*, table S4) of selective regimes between human groups. They are of particular relevance to understand how recent human demography might impact loci evolving under LTBS for millions of years or subsequent local adaptations through selective pressure shifts (e.g. de Filippo et al. 2016).

Our analyses indicate that LTBS in humans may be shaping variation in less than 2% of variable genomic positions, but that these on average overlap with about 8% of all protein-coding genes. Importantly, almost 70% of the candidate genes shared by at least same-continent populations cannot be ascribed to immune-related functions, suggesting that diverse biological functions, and the corresponding phenotypes, contain advantageous genetic diversity.

## Acknowledgements

We would like to dedicate this manuscript to Scott Williamson, in memoriam, for playing a fundamental role in the conception of NCD. We also thank Warren Kretszchmar for analyses on the properties of related statistics not included here, and Eric Green for his support of that work. We thank Michael DeGiorgio for assistance with BALLET, Felix Key with 1000 Genomes datasets, Michael Dannemann with the implementation of expression analyses, Stéphane Peyrégne for comments on the manuscript, David Reher, members of the Evolutionary Genetics Group (São Paulo), Alex Cagan and Svante Päabo for helpful comments. This work was supported by Fundação de Amparo à Pesquisa do Estado de São Paulo (grant numbers 11/12500-2 and 12/19563-2 to BDB and 12/18010-0 to DM) and the Max Planck Society (AMA).

